# Modeling glioblastoma invasion using human brain organoids and single-cell transcriptomics

**DOI:** 10.1101/630202

**Authors:** Teresa G Krieger, Stephan M Tirier, Jeongbin Park, Tanja Eisemann, Heike Peterziel, Peter Angel, Roland Eils, Christian Conrad

## Abstract

Glioblastoma multiforme (GBM) are devastating neoplasms with high invasive capacity. GBM has been difficult to study *in vitro*. Therapeutic progress is also limited by cellular heterogeneity within and between tumors. To address these challenges, we present an experimental model using human cerebral organoids as a scaffold for patient-derived glioblastoma cell invasion. By tissue clearing and confocal microscopy, we show that tumor cells within organoids extend a network of long microtubes, recapitulating the *in vivo* behavior of GBM. Single-cell RNA-seq of GBM cells before and after co-culture with organoid cells reveals transcriptional changes implicated in the invasion process that are coherent across patient samples, indicating that GBM cells reactively upregulate genes required for their dispersion. Functional therapeutic targets are identified by an *in silico* receptor-ligand pairing screen detecting potential interactions between GBM and organoid cells. Taken together, our model has proven useful for studying GBM invasion and transcriptional heterogeneity *in vitro*, with applications for both pharmacological screens and patient-specific treatment selection at a time scale amenable to clinical practice.

## Introduction

Glioblastoma (GBM) is the most frequent and most aggressive primary brain tumor^1,2^. Despite decades of intensive research, average survival time remains at 12-15 months from diagnosis^3^. Surgical resection of GBM tumors is rarely complete because the tumor aggressively infiltrates the brain, with cells interconnecting via long membrane protrusions (microtubes)^4^. The resulting network enables multicellular communication through microtube-associated gap junctions, and increases tumor resistance to cell ablation and radiotherapy^5^. Moreover, glioblastoma cells interact with normal brain cells via soluble factors or direct cell-cell contacts to promote tumor proliferation and invasion^6^.

Two major challenges have impeded progress in the development of new GBM therapies. Firstly, there is increasing evidence for substantial genetic^7,8^, epigenetic^9,10^ and transcriptional heterogeneity^11^ between and within human tumors. Recent advances in single-cell RNA sequencing (scRNA-seq) technology have enabled the transcriptomic analysis of numerous tumor entities at the level of individual cells. However, in the case of GBM, resection of primary samples has resulted in limited insight into interactions between infiltrating tumor and normal brain cells, as isolation of neoplastic cells from the tumor periphery has proven challenging^12^. How cellular heterogeneity of GBM cells and their interactions with normal brain cells relate to differences in proliferation or invasive capacity, which ultimately determine patient outcome, thus remains unknown.

A second challenge in the advancement of GBM therapies is the current lack of model systems to study defining properties of human GBM, especially invasion into the surrounding brain tissue. Previous in vitro models have suffered from limited physiological relevance or have been incompatible with the time scales for clinical decision-making^6^. Recent studies have shown that human cerebral organoids can be used as a platform for tumor cell transplantation or genetic engineering of tumors, enabling microscopic observation of tumor development^13–15^. However, tumor cell interactions with normal brain cells have not been addressed yet.

Here, we developed an experimental approach to study the interaction of GBM and normal brain cells of the neuronal lineage *in vitro*, on clinically relevant timescales of less than 4 weeks. We used iPSC-derived human cerebral organoids as a 3D scaffold for the invasion of patient-derived GBM cells and analyzed tumor microtube development by tissue clearing, confocal microscopy and semi-automated quantification. In addition, we performed scRNA-seq of GBM cells before and after co-culture with organoid cells and identified a transcriptional program induced by the interactions between tumor and normal brain cells, suggesting potential therapeutic targets.

## Results

### iPSC-derived cerebral organoids provide a scaffold for glioblastoma invasion

To study glioblastoma invasion in a physiologically relevant 3D context, we adapted an established protocol for human iPSC-derived cerebral organoid development^16^ to achieve streamlined and reproducible production of organoids. From 24 days of age, cerebral organoids were co-cultured with fluorescently labelled glioblastoma cells from four patient-derived cell lines (Fig. 1A and Supplementary Table 1). Samples were fixed after three days and subjected to tissue clearing using the FRUIT protocol^17^, enabling the visualization of tumor invasion by confocal microscopy. We found that tumor cells from all four GBM patients readily attached to and invaded into the organoids (Fig. 1B). Tumor cells formed protrusions reaching to other cells over short and long distances (Fig. 1B), consistent with tumor microtube formation observed *in vivo* in mice^4^. GBM cells primarily invaded into the neuronal layers of the organoids, with little invasion into neural progenitor rosettes (Fig. 1B).

**Figure 1.**
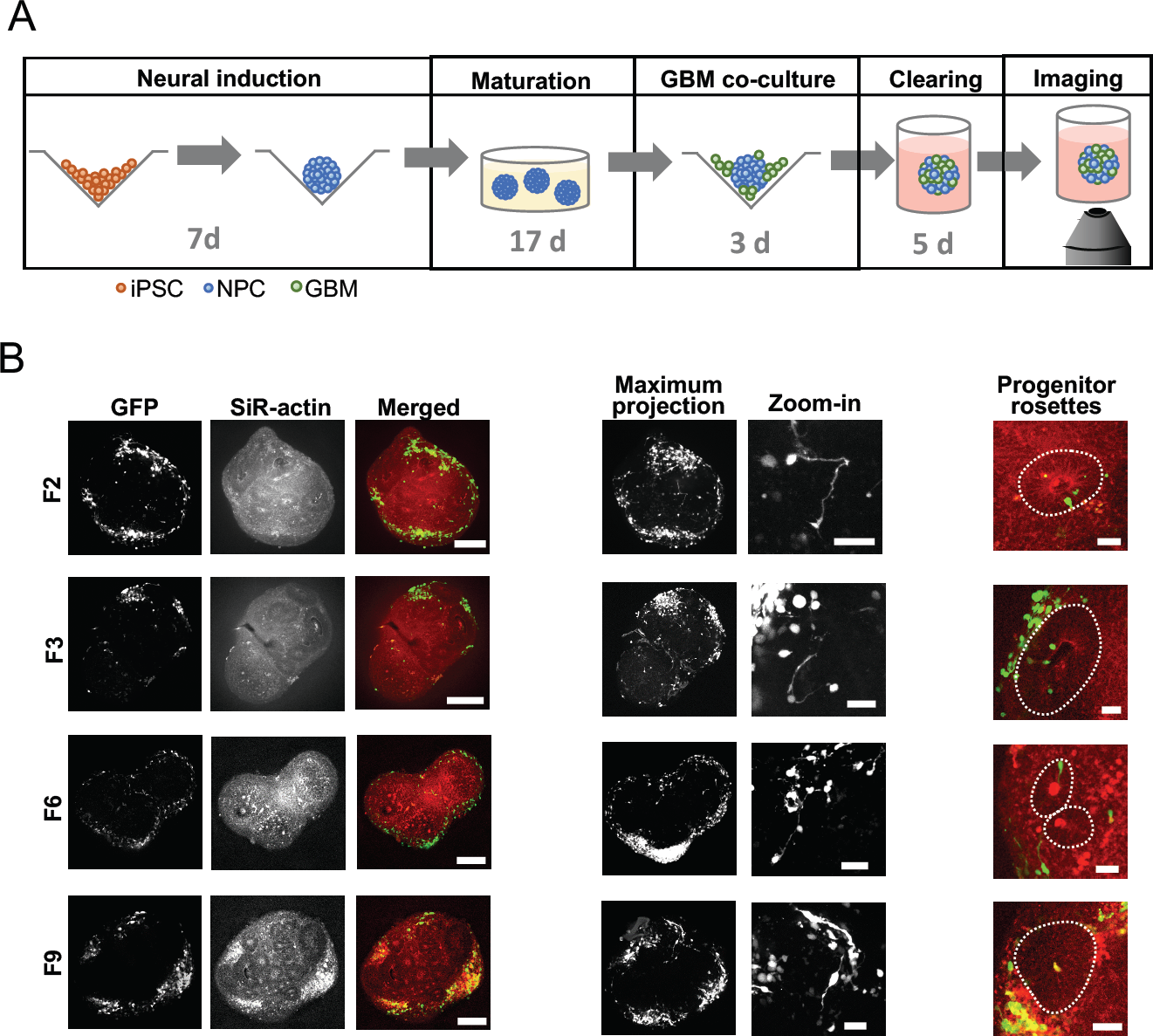
GBM invasion assay. **A)** Experimental protocol. Following 7 days of neural induction, organoids were transferred to Matrigel and matured for 17 days. Organoids were then enzymatically released and co-cultured with GFP-labelled GBM cells for 3 days. Samples were embedded in Matrigel again for fixation, tissue clearing and confocal imaging. **B)** GFP-labelled tumor cells from all four GBM patients invade into cerebral organoids (left; scale bars, 250 µm) where they form short-range and long-range connections (middle, maximum intensity projections over ∼200-250 µm depth; scale bars, 50 µm). Invasion is largely restricted to neuronal layers, outside of neural progenitor rosettes indicated by dotted lines (right; scale bars, 50 µm).

### Tumor microtube formation recapitulates *in vivo* behavior of GBM cells

We developed a semi-automated image processing workflow to analyze the invasion process quantitatively (Fig. 2A and Methods), which we applied to a total of 66 organoids (n=15-19 for each of the four patient GBM cell lines). We found that, for organoids of comparable sizes, the fraction of organoid volume taken up by tumor cells was similar across the four patient-derived cell lines (Fig. 2B and Supplementary Fig. 1A). The distribution of glioblastoma cells within organoids was assessed by calculating the distances between GFP^+^ voxels across the same set of organoids. Tumor cells spread widely in all cases (Supplementary Fig. 1B). To quantify invasion depth, we compared the distribution of distances of GFP^+^ voxels from the organoid surface across 12 similarly sized organoids for each patient-derived cell line. Invasion depths exceeded 100 µm in the majority of organoids (Fig. 2C), with some cells detected at approximately 300 µm from the organoid surface. While migration depth of the most invasive cells (90^th^ percentile of invasion depth) was uncorrelated with organoid size, we observed that cells from patients F6 and F9 were less invasive than cells from patients F2 and F3 (Fig. 2D and Supplementary Videos), suggesting that the *in vitro* model can reproduce intertumor heterogeneity in invasive behavior. By tracing membrane-bound cellular processes in images, we found that the number of microtubes per GBM cell ranged up to 6, with 2.2±0.1 microtubes on average (Fig. 2E). We quantified how many of these microtubes ended at other GBM cells, and identified between 0 and 4 such putative intratumoral connections per GBM cell, with an average of 1.2±0.1 connections (Supplementary Fig. 1C). Individual microtubes were up to 450 µm long (Fig. 2F). Consistent with our earlier observation of intertumoral heterogeneity of invasive capacity, we found that microtube lengths differed between cell lines (Fig. 2F). Interestingly, cell lines with higher invasive capacity also showed longer microtubes.

**Figure 2.**
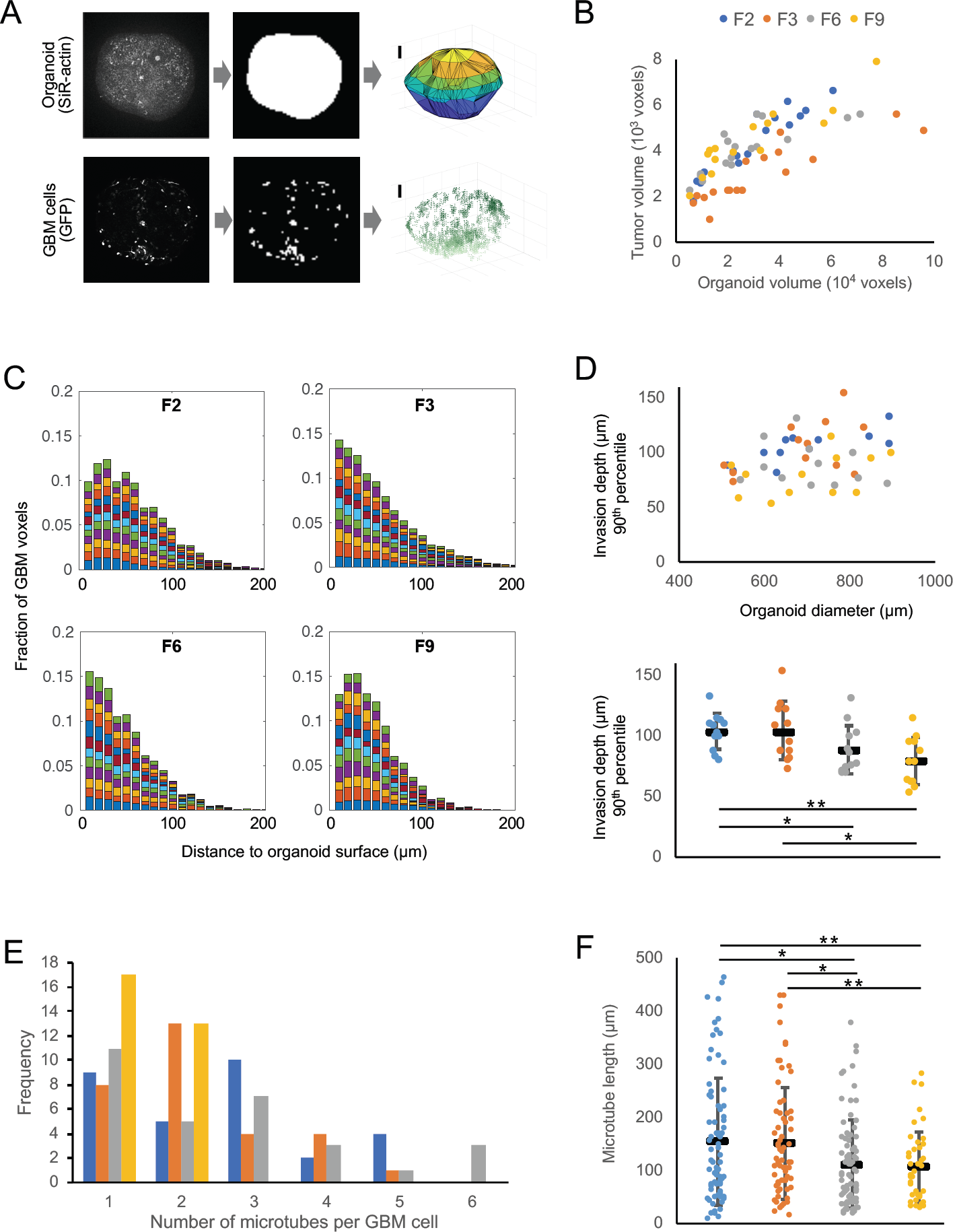
Morphological features of patient-derived GBM cells invading organoids. **A)** Image analysis workflow. To approximate the organoid surface, organoids were incubated with the live dye SiR-actin; following fixation and clearing, the actin signal was binarized and triangulated (top). Above-threshold GFP signal was used as a proxy for GBM cell location (bottom). Scale bars, 100 µm. **B)** Total tumor cell volume as a function of organoid volume. **C)** Distributions of distances of tumor voxels from the organoid surface of 12 organoids from each patient cell line with 500–900 µm diameter; each color represents one organoid. **D)** Invasion depth of the most invasive cells from each cell line (90^th^ percentile) compared to organoid size (top) and differences in invasion depth between patient cell lines (bottom; * p < 0.05, * * p < 0.01, two-sided Student’s t-test). **E)** Number of tumor microtubes per cell observed across 30 GBM cells from each patient. **F)** Microtube lengths ranged up to almost 450 µm, with GBM cells from patients F2 and F3 developing longer microtubes than cells from patients F6 and F9 (* p < 0.05, * * p < 0.01, two-sided Student’s t-test). In (D) and (F), black horizontal bars indicate mean values and error bars represent standard errors in the mean.

### scRNA-seq reveals transcriptional heterogeneity between tumors and after co-culture with organoid cells

Our imaging results confirm that iPSC-derived cerebral organoids represent an effective model system for quantifying GBM invasion and tumor microtube formation *in vitro*. To further study heterogeneity of and interactions between GBM and organoid cells at the transcriptome level, we developed a more efficient workflow that could be applied on clinically relevant timescales and at higher throughput. In the modified assay, dissociated 7-day-old cerebral organoids were mixed with GBM cells at a 1:1 ratio, and grown in co-culture for 3 days (Fig. 3A). GBM cells from all four patient-derived cell lines readily mixed with dissociated organoid cells (Fig. 3B and Supplementary Fig. 2A). With or without addition of GBM cells, dissociated organoid cells efficiently re-established the characteristic architecture of progenitor rosettes and neuronal layers observed in cerebral organoids, and membrane protrusions emanating from tumor cells were visible in all samples (Fig. 3C). After 3 days of co-culture, mixed spheroids were dissociated and subjected to scRNA-seq. For comparison, we also dissociated and sequenced recomposed spheroids of organoid cells that had not been mixed with GBM cells, referred to as neural progenitor cells (NPCs) below, and GBM cells from all four patient-derived cell lines that had been grown separately as spheroids in the same culture medium (Fig. 3A). Following pre-processing and quality control, we obtained 5,083 single-cell transcriptional profiles with approximately 1,400 genes detected per cell on average (Supplementary Fig. 2B). PCA-based clustering and 2D visualization by t-SNE maps revealed that GBM cells cultured alone clustered separately for each patient (clusters 5, 6, 7 and 9), confirming intertumoral heterogeneity (Fig. 3D). This was also highlighted by differential expression of putative marker genes for GBM subtypes^18^ across patient samples (Supplementary Fig. 2C). We further identified three clusters (clusters 0, 1 and 3) containing cells from the unmixed organoids as well as cells from all four mixed samples, and concluded that the latter represent the organoid cells in the mixed samples (Fig. 3D). The remaining clusters (clusters 2, 4 and 8) contain GBM cells from patients F2, F3 and F9 after co-culture with organoid cells. Note that as only six such cells were identified in the mixed sample from patient F6, they were excluded from further analyses and not displayed here.

**Figure 3.**
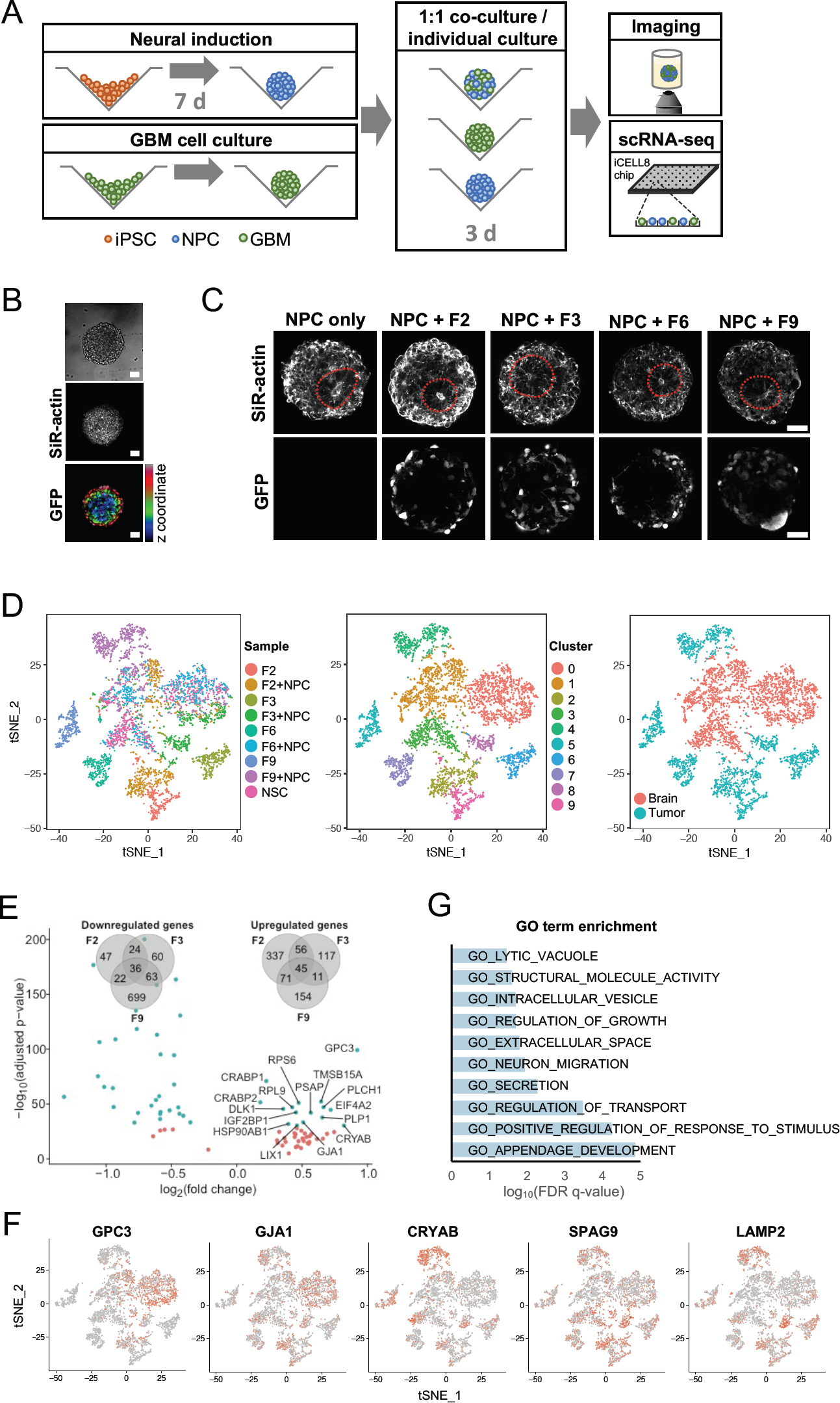
scRNA-seq analysis of GBM cell interactions with cerebral organoid cells. **A)** Protocol for the RNA-seq experiments. Following 7 days of neural induction, organoids and spheroids of lentivirally labelled GBM cells grown separately were enzymatically dissociated and mixed at a 1:1 ratio. After 3 days of co-culture, mixed spheroids were subjected to imaging or scRNA-seq using the iCell8 system. **B)** Tumor cells mixed efficiently with organoid cells (see also Supplementary Fig. 2). Scale bars, 50 µm. **C)** With or without addition of GBM cells, dissociated organoid cells re-established the characteristic 3D architecture of neural rosettes within 3 days. Scale bars, 50 µm. **D)** tSNE map showing all cells after QC and PCA-based clustering, colored by sample origin (left), by cluster (middle), and by organoid or tumor cell identity (right). In addition to three clusters containing organoid cells, GBM cells clustered separately for each patient and before or after co-culture with organoid cells. **E)** Volcano plot shows the 45 genes significantly up-or downregulated across all three patient-derived GBM cell lines upon co-culture with organoid cells (adjusted p-value < 0.05 for each patient separately). Venn diagrams quantify the overlap of differentially regulated genes detected from each patient. **F)** Expression of differentially regulated genes visualized on a tSNE map of all cells (tSNE representation identical to panel D). G) Gene Ontology (GO)-based gene set enrichment analysis of genes upregulated in all patient tumor cell lines upon co-culture with organoid cells.

### Mixing GBM and organoid cells leads to up-regulation of a shared set of genes across patients

Differential gene expression testing between GBM cell clusters from mixed and unmixed samples revealed hundreds of genes that were significantly up-or downregulated upon co-culture with organoid cells (adjusted p-value < 0.05, log(fold change) > 0.15), and an overlap of 45 genes that were upregulated in all patients (Fig. 3E). These included the homeobox transcription factor PAX6, normally expressed in forebrain neural stem cells; the gap junction protein alpha 1 (GJA1) coding for connexin-43, which connects tumor microtubes in GBM^4^; glypican-3 (GPC3), a cell surface heparan sulfate proteoglycan and Wnt activator whose expression correlates with invasiveness of hepatocellular carcinoma^19^; collagen COL4A5, an extracellular matrix constituent; and several lysosomal, vesicular and secretory proteins (Fig. 3F and Supplementary Table 2). Gene set enrichment analysis of the 45 coherently upregulated genes confirmed that genes relating to growth regulation, neuronal migration, extracellular secretion and stimulus response were enriched in this group (Fig. 3G). Our results thus show that interactions between GBM and organoid cells increase expression of genes required for GBM network formation and invasion.

### Potential ligand-receptor interactions between tumor cells and organoid cells

To investigate the nature of interactions between GBM and organoid cells, we considered the expression of 2,557 known ligand-receptor pairs^20^ across our samples, comprising a total of 1,398 unique genes. Of these, 317 genes were expressed in our data, with approximately 13% expressed differentially between the same cell types in unmixed and mixed samples (Fig. 4A). Calculating the number of potential interactions between brain and tumor cells based on the expression of complementary receptors and ligands, we detected substantial crosstalk between cell types (Fig. 4B). Hierarchical clustering of the number of cells potentially linked by each ligand-receptor pair revealed a group of ligand-receptor pairs that were expressed at low levels in the tumor-only and NPC-only cultures, but presented many potential interactions between tumor cells and organoid cells in the mixed cultures (Fig. 4C). These included several collagen-integrin interactions, glypican-3 binding to insulin-like growth factor 1 receptor (IGF1R) or the cell cycle regulator CD81, and non-canonical Notch signaling (DLK1/NOTCH1, DLK1/NOTCH2). Notably, despite the transcriptional heterogeneity we observed between patients, our approach detected consistently expressed potential interactions across all patient cell lines (Fig. 4C). Gene set enrichment analysis showed that the ligand-receptor pairs expressed at high levels in co-cultured samples are enriched for invasion-related genes (Fig. 4D). Specifically, putative interactions in which GBM cells present the ligand and NPCs the receptor are enriched for genes involved in neuron projection development and receptor binding, whereas ligand-receptor pairs communicating in the opposite direction are enriched for extracellular matrix proteins.

**Figure 4.**
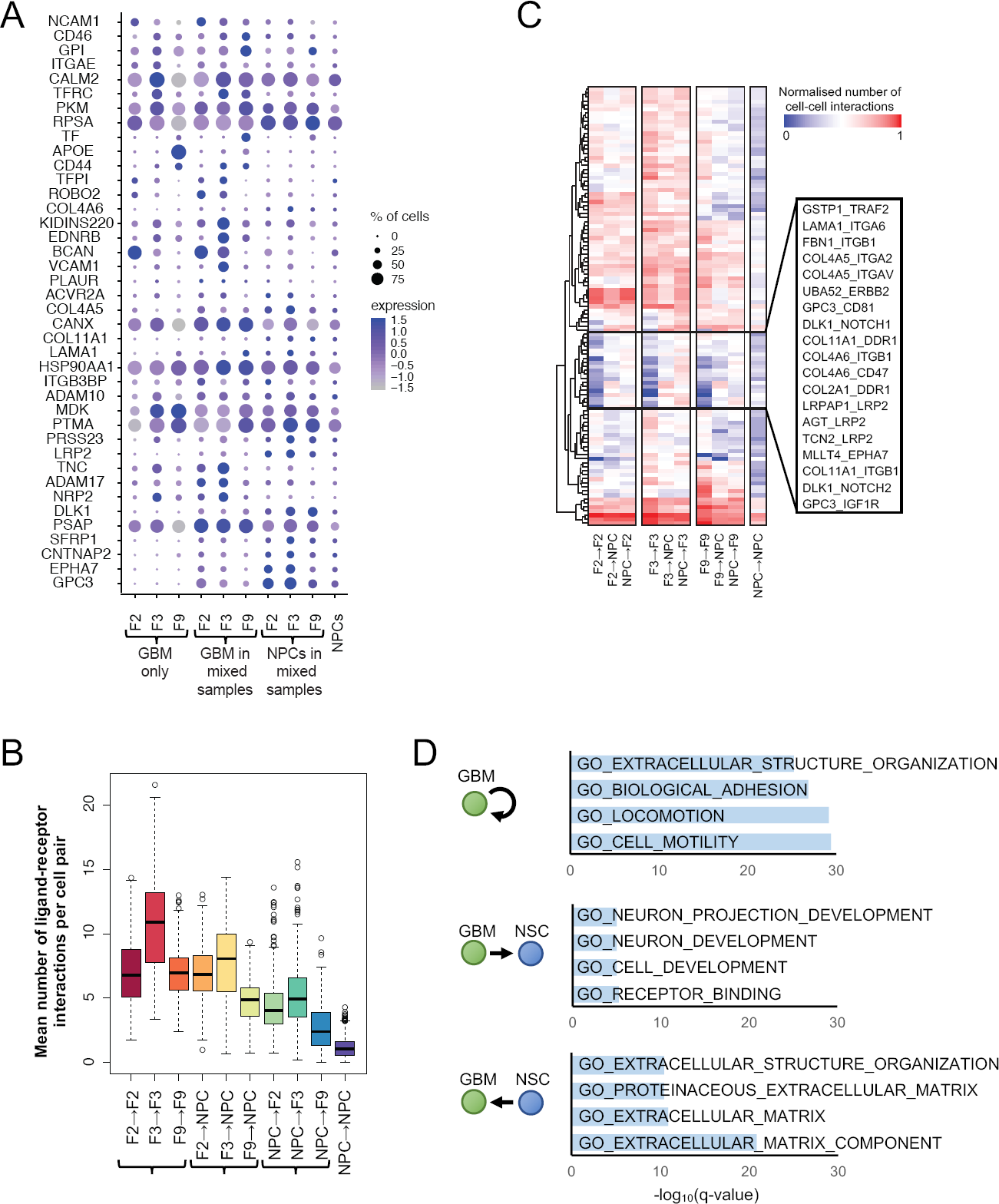
Potential ligand-receptor interactions between GBM and organoid cells. **A)** Expression of ligand and receptor genes that are significantly upregulated in NPCs or in at least one GBM cell line upon co-culture (adjusted p < 0.05), averaged across GBM-only samples (left), GBM cells or NPCs within mixed samples (middle), and the NPCs-only sample (right). Dot size corresponds to the fraction of cells expressing the gene in each group; dot color represents the average expression level. **B)** The mean number of ligand-receptor pairs potentially connecting cells pairs of the given cell types, based on RNA expression levels across GBM-only samples (left), GBM cells or NPCs within mixed samples (middle), and the NPCs-only sample (right). Center line, median; box limits, upper and lower quartiles; whiskers, 1.5x interquartile range; points, outliers. **C)** Mean number of cell-cell interactions for different cell type combinations and selected ligand-receptor pairs. Hierarchical clustering reveals a group of ligand-receptor pairs (middle box) that are expressed at low levels in GBM cells or NPCs alone, but are upregulated upon co-culture. **D)** Gene Ontology (GO)-based gene set enrichment analysis of ligand-receptor pairs preferentially expressed in the given cell type combinations.

## Discussion

Despite its enormous therapeutic and prognostic significance, efficient methods to characterize the process of GBM invasion into human brain at a quantitative or transcriptional level are currently lacking. In this study, we present an *in vitro* model system in which lentivirally labelled patient-derived GBM cells invade into human cerebral organoids. By tissue clearing and confocal imaging, our approach shows that tumor cells extend up to 450 µm long membrane-bound processes after three days of invasion, recapitulating the development of GBM microtubes that has been observed in resected primary tumors and replicated *in vivo* in mice^4^. Many of these processes terminate at distant tumor cells in our *in vitro* model, consistent with the development of an interconnected GBM network. By making GBM invasion experimentally accessible *in vitro* in a 3D tissue-like architecture, our experimental approach also enables the correlation of morphological phenotypes with transcriptional regulation by integration of imaging with single-cell sequencing. Here, scRNA-seq analysis of GBM and organoid cells separately or after co-culture revealed transcriptional changes induced by the interactions of tumor cells with their environment. Genes implicated in stimulus response, neuronal migration, secretion and extracellular matrix were coherently upregulated across all tumor samples when mixed with NPCs, indicating that GBM cells sense the presence of neuronal cells and reactively amplify the transcription of genes supporting their dispersion. Among the upregulated genes were GJA1 (coding for connexin 43), known to enable multicellular communication via gap junctions in GBM networks *in vivo*^4^, and GPC-3, which has received little attention in glioblastoma^21^ but whose expression evidentially correlates with higher invasive capacity in hepatocellular carcinoma^19^, where it also confers oncogenicity by activating the IGF signaling pathway through IGF1R^22^.

Heterogeneity between and within GBM tumors has impeded therapeutic progress for decades^23^, with no targeted therapy available yet^24^. Consistently, our imaging data shows that GBM cells from different patients varied in their invasive capacity. While additional patient samples and more single-cell transcriptional profiles would be necessary to robustly link intertumoral differences in invasion behavior with specific transcriptional changes, our results corroborated the high degree of transcriptional heterogeneity between patients^11^. However, we also detected a coherent element of transcriptional changes upon GBM and organoid cell co-culture indicating that targeting functional processes such as tumor microtube formation might improve therapeutic outcomes across patients. Our analysis of ligand-receptor pair expression identified candidate pairs that may contribute to the invasion process; further studies should explore the functional significance of these putative interactions.

The results presented here establish the biological relevance of our organoid-based experimental system and show that interactions between GBM and organoid cells result in transcriptional changes detected by scRNA-seq. In the future, we expect that our approach will further enable functional studies of GBM invasion that would be difficult or impossible to be conducted *in vivo*, including long-term imaging of network formation and multicellular communication. While we here used tissue clearing and confocal imaging for an end-point quantification of tumor invasion, GBM-invaded organoids are similarly amenable to live imaging by two-photon or light-sheet microscopy. By combining imaging with recent single-cell RNA sequencing methodologies that provide transcriptome data for greater cell numbers^25,26^, and with other single-cell sequencing modalities such as chromatin accessibility sequencing^27^, our model could thus help resolve the functional, transcriptional and epigenetic factors that are associated with different invasion behaviors of GBM or other tumors into human brain. It also provides the basis for high-content drug screens to assess patient-specific drug action on tumor and healthy brain cells, thus helping to identify the most effective drug at clinically relevant timescales.

## Acknowledgements

The authors would like to thank Katharina Jechow for technical laboratory support and Monika Langlotz for FACS assistance. TGK was partly supported by a DKTK postdoctoral fellowship from the Heidelberg School of Oncology. SMT and TE were supported by the PhD program of the Helmholtz International Graduate School for Cancer Research (DKFZ, Heidelberg).

## Author contributions

TGK, CC and RE conceived the study. TGK developed the protocol for GBM invasion into cerebral organoids, conducted experiments, analyzed the data and wrote the paper. SMT helped with scRNA-seq experiments and generated sequencing libraries. JP performed pre-processing of scRNA-seq data. TE, HP and PA provided GBM cells. All authors approved the manuscript.

## Methods

### GBM cell culture

Primary tumor samples were received from Frankfurt University Hospital (Edinger Institute). Informed consent was obtained prior to surgery. Experiments involving human patient material were performed in accordance with the Declaration of Helsinki and were approved by the ethics committee of the University Cancer Center Frankfurt, project number SNO_01_13. Patient-derived GBM cell cultures were established as described^28^. Cells were cultured in suspension culture in 75 cm^2^ ultra-low attachment flasks in Neurobasal medium (Gibco) supplemented with 2 mM L-glutamine (Life Technologies), 1x B27 (Gibco), 2 µg/ml heparin, 20 ng/ml EGF and 20 ng/ml bFGF (R&D Systems). For passaging, spheroid cultures were dissociated using Accutase (StemCell Technologies) when they had reached diameters of ∼100 µm, every 1-3 weeks.

### Lentiviral labelling and FACS sorting

Second-generation replication-incompetent lentivirus was produced by FuGENE transfection (Promega) of HEK293T cells with the expression plasmid LeGO-G2, complemented with the packaging plasmids psPAX2 and pMD2.G (all from Addgene). GBM cells were infected on three consecutive days by spinoculation at 800 g for 30-60 minutes, and GFP-expressing cells were isolated by FACS. Lentivirally labelled GBM cells were maintained in neural maintenance medium and passaged every 1-3 weeks as described above.

### Organoid culture

The iPS cell line 409b2 was obtained from Riken Institute (Wako, Japan). For routine culture, iPSCs were maintained in mTeSR1 medium (StemCell Technologies) on tissue culture plates coated with Matrigel (Corning) and passaged every 4-5 days using Gentle Cell Dissociation Reagent (StemCell Technologies). For organoid seeding, iPSCs were dissociated with Accutase (StemCell Technologies) and transferred into AggreWell plates in Neural Induction Medium (StemCell Technologies) with 10 µM Y-27632 (StemCell Technologies) at a density of 1,000 cells per cavity, following manufacturers’ instructions. Spheroid formation was confirmed visually after 24 hours, and spheroids were maintained in Neural Induction Medium (StemCell Technologies) with daily medium changes. After 5 days, spheroids were harvested from the AggreWell plates and embedded in Matrigel. Medium was changed to neural maintenance medium, a 1:1 mixture of N-2 and B-27-containing media (N-2 medium: DMEM/F-12 GlutaMAX, 1× N-2, 5 µg/ml insulin, 1 mM L-glutamine, 100 µm nonessential amino acids, 100 µM 2-mercaptoethanol; B-27 medium: Neurobasal, 1× B-27, 200 mM L-glutamine) supplemented with 50 U/ml penicillin and 50 mg/ml streptomycin, and exchanged every 2 days.

### Organoid invasion assays

On day 24 of culture, organoids were removed from Matrigel by incubation with Dispase (Sigma) at 37°C, and transferred to individual wells of a GravityTRAP ULA Plate (PerkinElmer). Labelled GBM cells were dissociated with Accutase, and added to the organoid plate in neural maintenance medium at a concentration of 1,000 cells per well. Plates were centrifuged at 100 g for 3 min before returning to the incubator. After 2 days, Organoids were live stained with 100 nM SiR-actin (Spirochrome). The following day, organoids were harvested, fixed in 2% PFA for 30 min, and embedded in Matrigel for immobilization. Tissue clearing was performed following the FRUIT protocol as described^17^. Confocal images were acquired on a LSM780 Axio Observer confocal laser scanning microscope (Zeiss).

### Image analysis

Image pre-processing was performed in ImageJ as follows: To obtain a representation of the organoid, the SiR-actin signal was subjected to brightness adjustment and Gaussian blurring (sigma = 2), followed by scaling to 2912 µm^3^/voxel by bicubic interpolation, and binarization using the Triangle method. To identify the voxels occupied by GBM cells, the GFP signal was subjected to brightness adjustment and Rolling-Ball background subtraction (radius = 64), followed by scaling to 2912 µm^3^/voxel by bicubic interpolation, and binarization using Otsu’s method.

Downstream image processing was performed in MATLAB as follows: Using binarized images, single cells in the Matrigel surrounding organoids were excluded by connected component analysis using the ‘bwlabeln’ function, and holes inside the organoid were filled with the ‘imfill’ function. The organoid surface was approximated by Delauney triangulation using the ‘delaunayTriangulation’, and normal distances from GBM-occupied voxels to the organoid surface were calculated with the ‘point2trimesh’ function. We performed hierarchical clustering of voxels based on Euclidean distances and calculated distance matrices to visualize dispersion of GBM cells within organoids.

Processes were tracked using the Simple Neurite Tracer in ImageJ. A total of 120 cells (n=10 cells each from n=3 organoids for each patient) were randomly selected, and all their processes tracked.

### scRNA-seq

#### Sample preparation

For scRNA-seq of GBM cell lines, cells were cultured in neural maintenance medium for 1-3 weeks after passaging, until the largest spheroids were ∼100 µm in diameter. Cells were then dissociated with Accutase (StemCell Technologies), washed twice in PBS, and passed through a 20 µm cell strainer (PluriSelect). For scRNA-seq of co-cultured GBM and organoid cells, NPC spheroids were generated by inducing iPSCs AggreWell plates as described above. After 7 days, NPC spheroids and lentivirally labelled GBM cells were dissociated using Accutase (StemCell Technologies), mixed in a 1:1 ratio, and replated onto AggreWell plates at 1,000 cells per cavity in neural maintenance medium. After 3 days, mixed spheroids were dissociated using Accutase (StemCell Technologies), washed twice in PBS, and passed through a 20 µm cell strainer (PluriSelect).

#### Cell isolation and library preparation

Single-cell suspensions were stained with Hoechst and Propidium Iodide (ReadyProbe Cell Viability Imaging Kit, Invitrogen) for 10 min at room temperature and cell numbers and viability were checked with a Countess automated cell counter (Thermo Fisher). Samples were discarded if cell viability was below 85%. The TakaraBio iCELL8 system and the associated Rapid Development Protocol (in-chip RT-PCR amplification) were used for single cell isolation, reverse transcription and cDNA amplification^29^. Briefly, cell suspensions were distributed into a nanowell chip containing oligo-dT primers with a unique barcode for every well. Chips were imaged using an automated fluorescence microscope and frozen at −80°C until further use. Nanowells occupied by single cells were identified using the CellSelect software and manually curated in order to exclude non-detected doublets or dead cells. After thawing frozen chips, 50 nl of RT/Amp solution was dispensed into selected nanowells (Master mix: 56 µl 5 M Betaine, 24 µl 25 mM dNTP mix (TakaraBio), 3.2 µl 1 M MgCl_2_ (Invitrogen), 8.8 µl 100 mM Dithiothreitol (TakaraBio), 61.9 µl 5x SMARTScribe™ first-strand buffer, 33.3 µl 2x SeqAmp™ PCR buffer, 4.0 µl 100 µM RT E5 Oligo, 8.8 µl 10 µM Amp primer (all TakaraBio), 1.6 µl 100% Triton X-100 (Acros), 28.8 µl SMARTScribe™ Reverse Transcriptase, 9.6 µl SeqAmp™ DNA Polymerase (TakaraBio)). After in-chip RT/Amp amplification (18 amplification cycles, in-chip RT/Amp Rapid Development protocol) inside a modified SmartChip Cycler (Bio-Rad), libraries were pooled, concentrated (DNA Clean and Concentrator−5 kit, Zymo Research) and purified using 0.6x Ampure XP beads. Concentration and quality of cDNA was assessed by a fluorometer (Qubit) and by electrophoresis (Agilent Bioanalyzer high sensitivity DNA chips). Next generation sequencing libraries were constructed using the Nextera XT kit (Illumina) following the manufacturer’s instructions. Final libraries were sequenced with the NextSeq 500 system in high-output mode (paired-end, 21 × 70 for v1, 24 × 67 for v2 chip).

### scRNA-seq data analysis

#### Pre-processing, quality control and normalization

For pre-processing of single-cell RNA-seq data, an automated in-house workflow based on Roddy (https://github.com/TheRoddyWMS/Roddy) was used. Read quality was evaluated using FastQC. iCELL8 library barcodes from the first 21 bp reads were assigned to the associated nanowell with the Je demultiplexing suite^30^. Remaining primer sequences, Poly-A/T tails and low-quality ends (<25) were trimmed using Cutadapt. Furthermore, since NextSeq (Illumina) encodes undetected bases as incorrect ‘Gs’ with high quality, Cutadapt’s ‘—nextseq-trim’ option was used for improved quality trimming. Trimmed reads were mapped to the reference genome hs37d5 (derived from the 1000 genomes project) using the STAR aligner. Mapped BAM files were quantified using featureCounts with reference annotation gencode v19.

RNA-seq libraries that contained less than 150 detected genes or more than 15% mitochondrial reads were filtered out. Adapting a previously published approach^31^, aggregate expression for each gene across all cells was calculated as *E*_*a*_ = log(mean*[E*_*j,1*…*n*_] + 1), where *E*_*j*_ is the counts-per-million expression value of the gene in cell *j*. 8,533 genes with *E*_*a*_ > 2 were retained for analysis.

#### Comparison of tumor cells from individual and co-cultured samples

Filtered and normalized data of all patients was combined to identify NPCs and tumor cells in each sample. Using the Seurat package as implemented in R^26^, principal component analysis (PCA) was performed prior to clustering, and the ‘FindClusters’ function (with resolution = 0.4) was run on the first 9 principal components only. Results were visualized by tSNE^32^. Clusters containing cells from the NPC-only sample were identified as ‘brain’, whereas clusters containing only GBM cells were identified as ‘tumor’. After manual splitting of one of the resulting clusters, we obtained 10 clusters representing brain cells (3 clusters), tumor cells from co-cultured samples, or tumor cells from unmixed samples. Differential expression between mixed and unmixed tumor cells was evaluated using the ‘FindMarkers’ function in Seurat. Gene set enrichment analysis^33^ was performed by computing overlaps between identified gene signatures and Gene Ontology (GO_C5) gene sets derived from the Molecular Signature Database (MSigDB, https://software.broadinstitute.org/gsea/msigdb).

#### Analysis of ligand-receptor interactions

Potential receptor–ligand pairings were analyzed using 2,557 previously published receptor–ligand pairs consisting of 1,398 unique genes^20^, of which 317 were expressed in our data. Adapting a previously published approach^34^, we constructed a cell-cell interaction matrix by summing for each pair of cells from the same sample the number of ligand-receptor pairs potentially connecting the pair, with one cell type expressing the receptor and the other the ligand (normalized expression cutoff >0.5). To identify ligand-receptor interactions with likely significance for the invasion process, we then considered each ligand-receptor pair in turn, and calculated the number of cell pairs connected by this ligand-receptor interaction for each possible cell type combination (tumor–tumor, brain–brain, tumor–brain), for each sample. The resulting interaction matrix was normalized to the maximum possible number of cell-cell interactions. To identify ligand-receptor pairs with coherent differential expression across patients, we considered only those ligand-receptor pairs with mean normalized expression greater than 0.5 times the mean across all pairs for all tumor-only samples, all pairings from mixed samples where tumor cells express the ligand, all interactions from mixed samples where tumor cells express the receptor, or the NPC-only sample. These interactions were clustered based on complete linkage of Euclidean distances and visualized using the heatmap.2 package in R. For Gene Ontology (GO) analysis, we considered those ligand-receptor pairs with mean normalized expression greater than 0.5 times the mean across all pairs for one cell type interaction (GBM→NPC, NPC→GBM, GBM→GBM, NPC→NPC), and less than 0.5 times the mean for the other three. No putative GBM→GBM interactions fulfilled these criteria.

## Data availability

Raw sequencing data has been deposited at the European Genome-Phenome Archive (http://www.ebi.ac.uk/ega/) under accession number EGA#XXXXXX.

## Code availability

Scripts used for analyzing transcriptome data and image data (in R, Fiji and MATLAB) are available from the authors upon request.

