## Supplementary Figures and Tables for "Modeling glioblastoma invasion using human brain organoids and single-cell transcriptomics"

### *Supplementary Figure 1:*

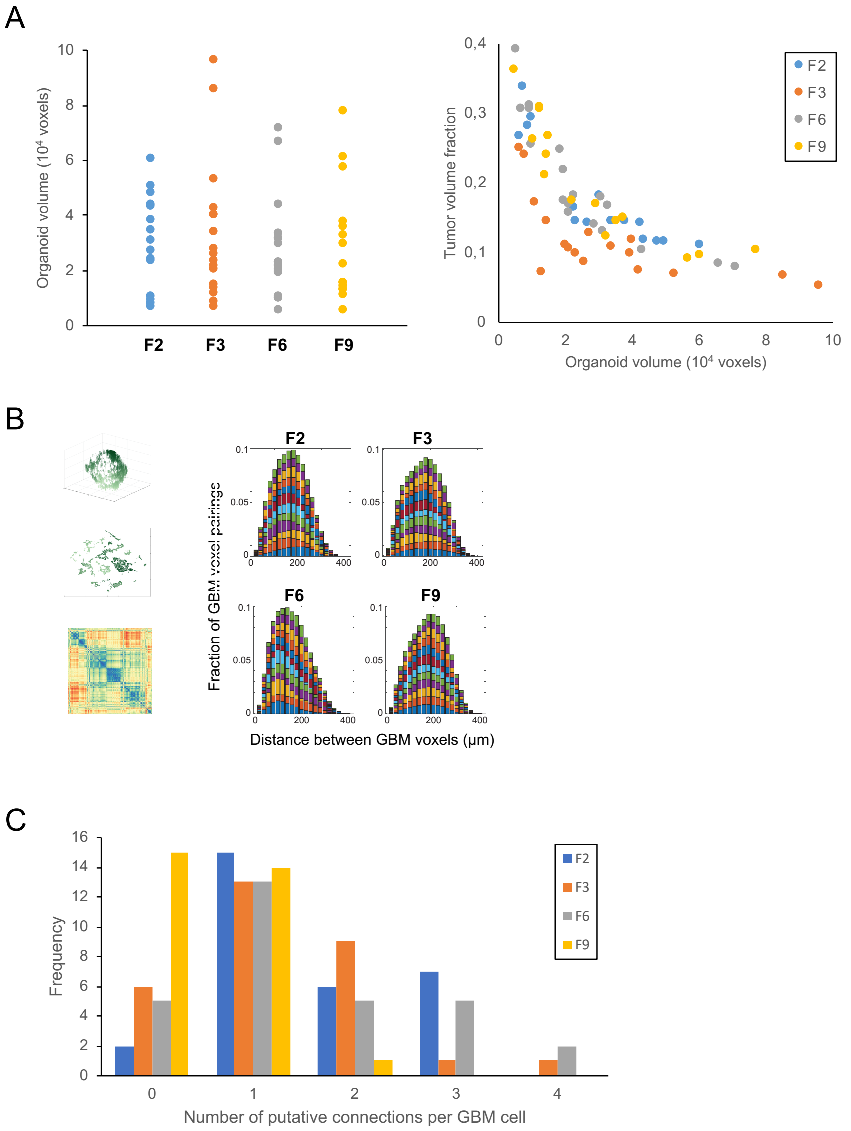


A) Organoids of similar sizes were selected for all four patient-derived GBM cell lines for experiments and quantification (left). A similar fraction of organoid volume was invaded by GBM cells from all four patient-derived cell lines (right). B) The distribution of distances between tumor voxels within organoids was similar for all four patient-derived cell lines. For visualization, a representative 3D rendering of GBM voxels is depicted (top left) along with a tSNE projection (below) and the corresponding distance matrix (bottom left). The distribution of distances between tumor voxels is shown for 12 representative organoids for each patient-derived cell line, with 500-900 µm diameter (right). C) Putative intratumoral connections, i.e. microtubes ending at other tumor cells, ranged in number from 0 to 4 per GBM cell.

*Supplementary Figure 2:*

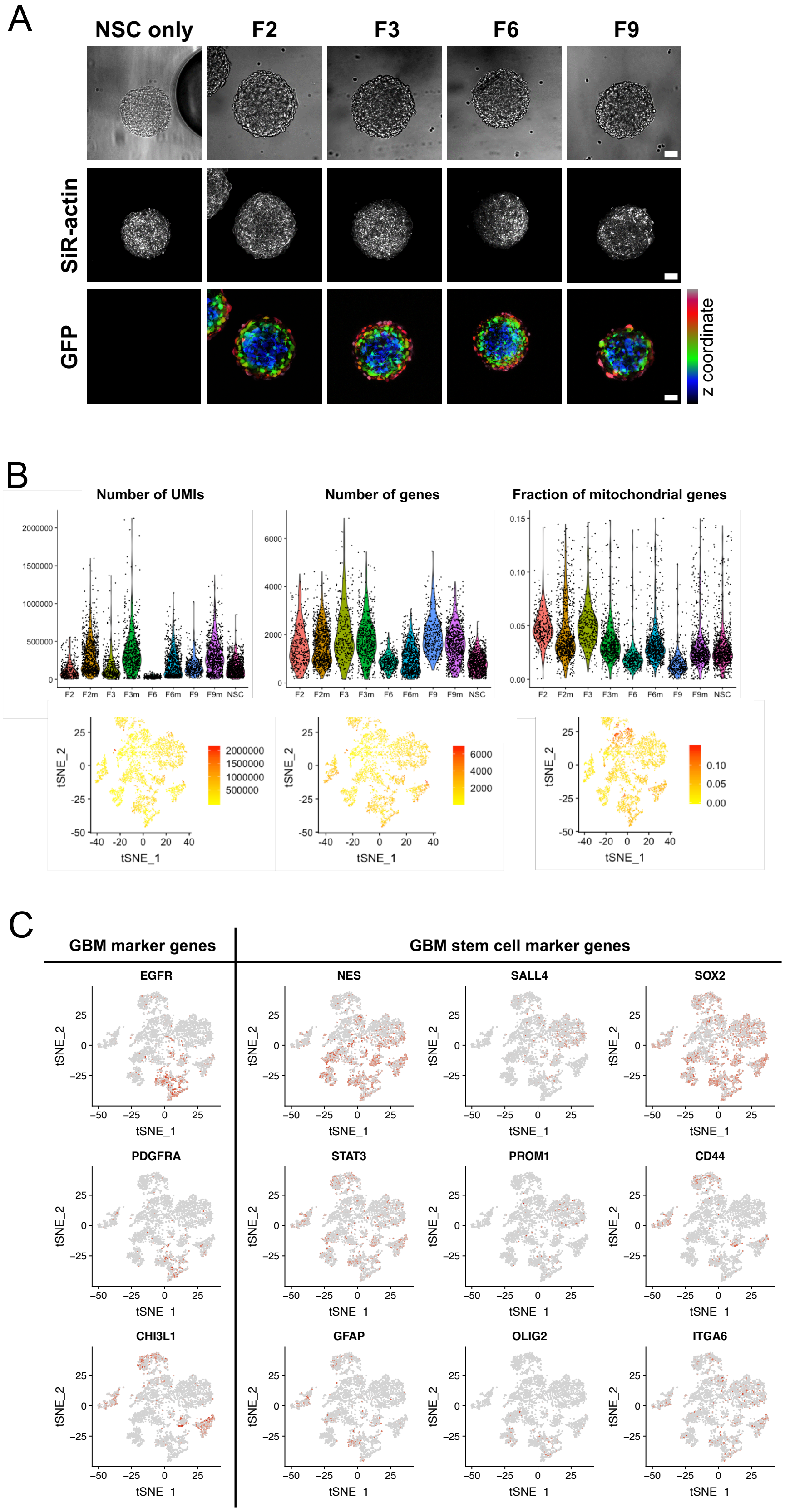


A) Tumor cells from all patient cell lines mix efficiently with organoid cells. Scale bars, 50 µm. B) Top row shows the number of UMIs, the number of genes, and the percentage of mitochondrial genes in each sample. Bottom row shows the same variables visualized on a tSNE map of all cells (identical to the tSNE representation shown in Fig. 3D). C) Expression of putative GBM marker genes (left) and putative GBM stem cell marker genes (right) visualized on a tSNE map (identical to the tSNE representation shown in Fig. 3D; grey, low expression; red, high expression).

### *Supplementary Table 1:*

| Cell line | Classification | Primary/ Recurrence | Sex | Age | Localization | Treatment | IDH[-] | ATRX[+] | MGMT promoter methylation |
|---|---|---|---|---|---|---|---|---|---|
| GBM-F2 | Glioblastoma, WHO grade IV | Primary tumor | M | 65 | left occipital | none | ✓ | ✓ | ✓ |
| GBM-F3 | Glioblastoma, WHO grade IV | Recurrence (after resection and re-resection) | M | 52 | right occipital | Radio-chemotherapy and surgery | ✓ | ✓ | unknown |
| GBM-F6 | Glioblastoma, WHO grade IV | Primary tumor | M | 68 | right temporo-parietal | none | ✓ | ✓ | ✓ |
| GBM-F9 | Glioblastoma, WHO grade IV (giant cell component) | Primary tumor | F | 71 | right temporal | none | ✓ | ✓ | ✓ |

### *Supplementary Table 2:*

Genes coherently upregulated in GBM lines upon co-culture with organoid cells

GPC3
CRABP1
CRABP2
PLP1
PLCH1
ARGLU1
TMOD3
RPS6
EIF4A2
RPL9
IGF2BP1
ARRDC3
SCG5
PSAP
TMSB15A
SPAG9
DLK1
LIX1
ANKRD36
SUGP2
PRTG
HSPA8
POLR2B
ANKRD36C
MORF4L2
EIF2S3
POLA1
PUM1
NKTR
VPS13B
DDX3X
COPA
LARS
VPS41
PAX6
HSP90AB1
PRUNE2
GNPDA1
HSP90AA1
CRYAB
BLOC1S5
GJA1
COL4A5
LAMP2
KDM4A

### *Supplementary Videos 1-4:*

Representative example z-scans of cleared organoids after GBM invasion from the four patient cell lines (1: F2, 2: F3, 3: F6, 4: F9).
